# Bi-phasic patterns of age-related differences in dopamine D1 receptors across the adult lifespan

**DOI:** 10.1101/2022.05.24.493225

**Authors:** Jarkko Johansson, Kristin Nordin, Robin Pedersen, Nina Karalija, Goran Papenberg, Micael Andersson, Saana M. Korkki, Katrine Riklund, Marc Guitart-Masip, Anna Rieckmann, Lars Bäckman, Lars Nyberg, Alireza Salami

**Affiliations:** Department of Radiation Sciences, Diagnostic Radiology, Umeå University, S-90187 Umeå, Sweden; Umeå Center for Functional Brain Imaging (UFBI), Umeå University, S-90187 Umeå, Sweden; Aging Research Center, Karolinska Institutet & Stockholm University, Tomtebodavägen 18A, S-17165, Stockholm, Sweden; Department of Integrative Medical Biology, Umeå University, S-90187 Umeå, Sweden; Wallenberg Center for Molecular Medicine, Umeå University, Umeå, Sweden; Max Planck UCL Centre for Computational Psychiatry and Ageing Research, University College London, London, UK; The Munich Center for the Economics of Aging, Max Planck Institute for Social Law and Social Policy, 80799 Munich, Germany

## Abstract

The dopamine (DA) system, particularly D1-like DA receptors (D1DR), declines across the adult life. The functional consequences of reduced D1DR has been hypothesized to vary across life periods, but the precise timing of these periods is unknown. To examine distinct phases in age-related D1DR reductions, we studied 180 healthy adults (90 females, 20-80 years), who underwent D1DR PET assessment using [^11^C]SCH23390. A bi-phasic pattern of age-related D1DR differences was revealed, with an inflection point at approximately 40 years of age. Notably, D1DR levels before and after the inflection showed opposing relations to neurocognitive functions, in concordance with distinct consequences of D1DR differences during development and in old age. Furthermore, D1DR reductions in later life were linked to age-related cerebrovascular consequences. These results support a distinction between D1DR reductions in early adulthood from those later in life, and suggest less dramatic and more malleable DA losses in aging than previously suggested.

## Introduction

The dopamine (DA) system undergoes profound changes across the lifespan, along with concomitant alterations in cognition ((Bäckman et al., 2010; Islam et al., 2021; Li et al., 2010; Reynolds and Flores, 2021a; Wahlstrom et al., 2010), for reviews). In childhood and adolescence, ongoing maturation of the prefrontal DA system is associated with excessive DA activity (Islam et al., 2021; Reynolds and Flores, 2021a; Wahlstrom et al., 2010, 2007), constraining the development of executive functions (Diamond, 2009; Diamond et al., 2004; Diamond and Baddeley, 1996; Klune et al., 2021). At older ages, DA losses may result in insufficient DA modulation (Lindenberger et al., 2008; Nagel et al., 2008), and decline in multiple cognitive domains (Bäckman et al., 2000, 2006; Karalija et al., 2019; Nyberg et al., 2010; Volkow et al., 1998). Consequently, characterization of age-related DA trajectories holds promise to illuminate the neural underpinnings of cognitive alterations across the lifespan.

The DA D1- and D2-like receptors (D1DR, D2DR) and dopamine transporters (DAT) reduce with advancing age, typically following a linear pattern (see (Karrer et al., 2017) for a review). However, the majority of past investigations are based on extreme age group comparisons precluding precise characterization of age-related trajectories. In this regard, D1DRs are of prime interest since they are the most abundantly expressed DA receptor subtype in the brain (Hall et al., 1994; Jaber et al., 1996), strongly implicated in the maturation of the PFC (Reynolds and Flores, 2021b), and cognition (Williams and Goldman-Rakic, 1995), and may exhibit more protracted development compared to the D2DRs (Rothmond et al., 2012; Weickert et al., 2007). I*n vivo* and *post mortem* studies suggest negative age differences for D1DR across the adult lifespan, with marked reductions already in young adulthood (∼20 – 40 y; (Rinne et al., 1990; Seeman et al., 1987), see Karrer et al. 2017 for a review). In addition, an imaging study including juvenile participants found a continuous trajectory of D1DR reductions across the age span of 10 – 32 y (Jucaite et al., 2010). These findings, together with indications of D1DR expression peaking in the third decade of life (Rothmond et al., 2012; Weickert et al., 2007), suggest protracted D1DR development during early adulthood. However, at present, there is a paucity of well-powered lifespan D1DR studies, thus preventing firm conclusions regarding different types of reduction, their heterogeneity across brain regions, and their relation to measures of brain health, function and cognition.

D1DR development beyond adolescence has implications for adult cognition. An inverted-U shaped model of differences in DA modulation across the lifespan (e.g. (Li et al., 2010)) posits compromised cognitive performance during development due to *excessive* DA modulation, whereas in later ages poorer cognition couples with insufficient DA. Hence, if protracted D1DR development manifests in adult age, the model posits a coupling between high D1DR expression and markers of excessive DA modulation and poorer cognition in young. DA is a critical player to enhance neural signal-to-noise ratio (Cools and D’Esposito, 2011; Frank and Fossella, 2011; Goldman-Rakic et al., 2000; Li and Sikström, 2002), and thus, age-related DA differences likely encompass alterations in neural communication (Garzón et al., 2021; Geerligs et al., 2015; Goh, 2011; Park et al., 2004; Pedersen et al., 2021). Functional connectivity (FC) in the striatum – the most DA-rich region in the brain – might hence serve as an *in-vivo* marker of altered DA modulation. Past studies have linked striatal FC changes to DA maturation in adolescence (Parr et al., 2021), whereas age-related changes in striatal FC was negatively linked to memory in older adults (Ystad et al., 2010), possibly reflecting variations in DA integrity (Nyberg et al., 2016; Rieckmann et al., 2018).

Here, we use the largest D1DR data set to date, the population-based DyNAMiC sample covering adult age span (n=180, 20 – 80 y, 30 per decade, 90 females;(Nordin et al., 2022)), to characterize break points in age-related D1DR trajectories, critical ages of transition between potential distinctive periods, and explore their relations to cerebrovascular brain integrity, brain function and cognition. Two alternative age-related trajectories are plausible in view of past findings. Firstly, a linear pattern has been most frequently reported in past studies of D2DR, DAT (Karrer et al., 2017), and D1DRs (Rinne et al., 1990; Seeman et al., 1987; Suhara et al., 1991). By a linear account, a single neurophysiological mechanism would continuosly exert a negative influence on D1DR expression across the entire adult lifespan. Secondly, an alternative view is a non-monotonous pattern, implicating multiple neurophysiological factors that have distinct influences across the adult lifespan. A similar model has been suggested for brain morphology (Fjell et al., 2013; Sowell et al., 2003). If a non-monotonous pattern emerges, brain-wide reductions in early adulthood, followed by attenuated rates of differences might be observed if D1DR development wanes during early adulthood (c.f. Rothmond et al., 2012; Weickert et al., 2007). Alternatively, D1DRs may display preservation across most of adulthood followed by accelerated and regionally variant rates of decline in older age. We predict that D1DR reductions in older age, in particular in the striatum (Iadecola, 2013), may be attributed to cerebrovascular integrity as assessed using white matter lesion volumes (Karalija et al., 2019; Rieckmann et al., 2016). We further predict that high D1DR couples with impaired striatal FC and poorer cognition during development, whereas in senescence low D1DR associates with poorer cognition and aberrant striatal FC.

## Results

### Spatial distribution of D1DRs across the brain

The spatial distribution of D1DR availability followed the patterns reported previously (Ito et al., 2008): D1DR binding was highest in striatal regions, followed by frontal, temporal, and parietal regions (Fig 1). Cortical D1DR binding was approximately five times lower than in the striatum.

**Figure 1:**
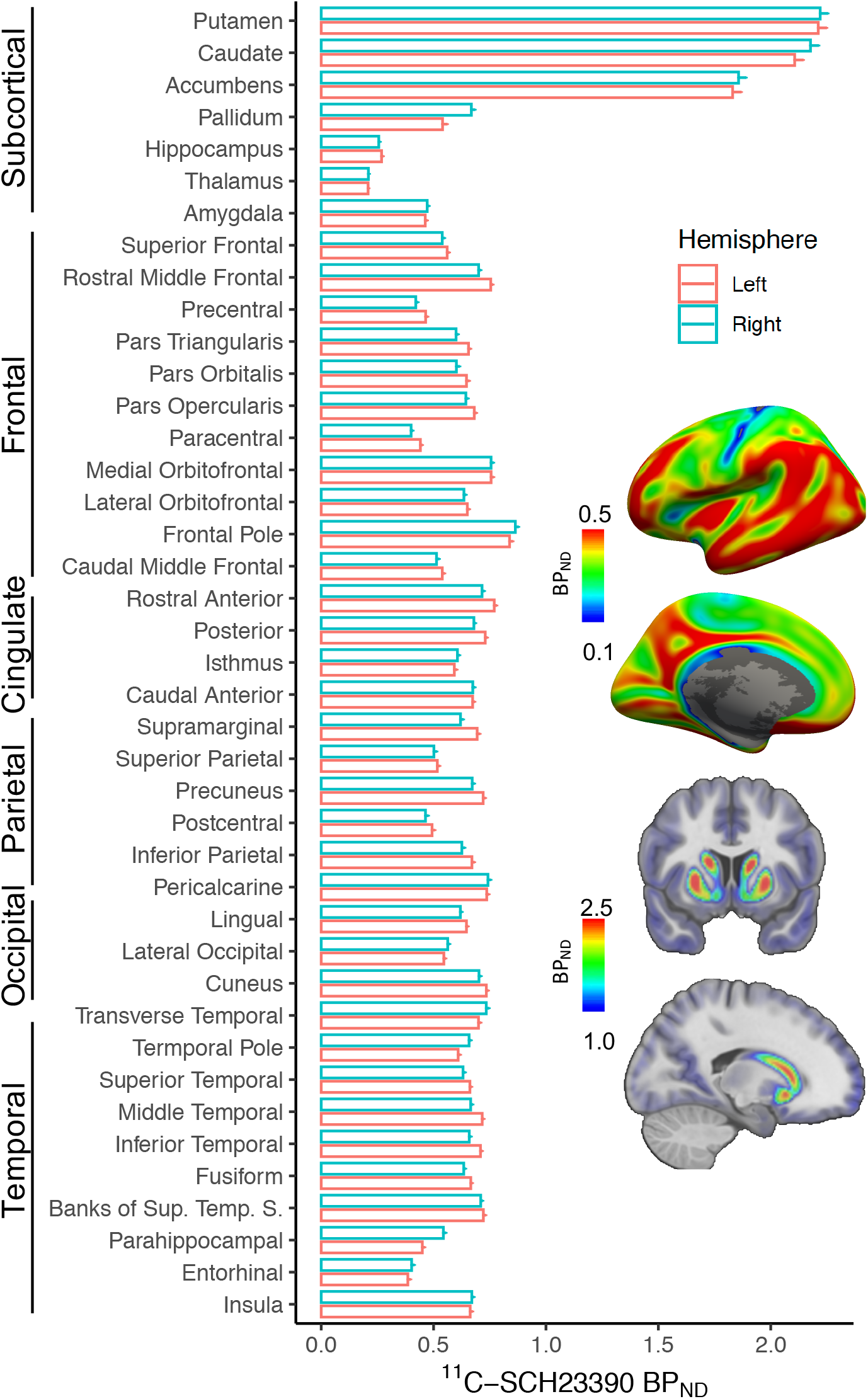
Average [^11^C]SCH23390 BP_ND_ in early adulthood (age < 40 y, n=60). Bar-graph: mean(+se) in Desikan-Killiany-based regions-of-interest in both hemispheres. Images: distribution of surface-mapped PVE-corrected cortical (top) and volumetric subcortical (bottom) [^11^C]SCH23390 BP_ND_.

### Age sensitivity of D1DRs across the brain

Age effects were examined within automatically defined anatomical regions-of-interest (ROI) according to Desikan-Killiany atlas and subcortical structures (Desikan et al., 2006; Fischl, 2012), providing whole-brain coverage. ROI-based analysis (Supplementary Fig 1) revealed significant negative age-related D1DR differences in 87% of the ROIs (Bonferroni-corrected ps < 0.05, adjusted r^2^s = 0.12 – 0.63, mean ± sd = 0.36 ± 0.12, % BP difference per decade = [-8.67, -1.81] %, mean ± sd = -4.58 ± 2.16, Supplementary Table 1), and no systematic hemispheric lateralization of the age effects (paired t-test *p* = 0.45, Pearson’s r = 0.8, Supplementary Figure 2). These results suggest wide-spread age-related reductions to D1DRs across the adult life span.

### Bi-phasic trajectories of D1DRs across the adult life span

To characterize age-related D1DR trajectories striatal ROIs were distinguished from cortical ROIs, owing to potential differences between nigrostriatal and mesocortical DA pathways, as well as between DA-rich striatal versus less densely DA-innervated extrastriatal regions. In a data reduction step, hierarchical GAM was run on the cortical ROIs in order to reduce the number of ROIs exhibiting mutually similar relationships with age (see Supplemental Material). This analysis identified four composite cortical ROIs (Fig 2a, Supplemental Table 3): prefrontal cortex (PFC), operculum & anterior cingulate cortex (ACC), and posterior and central cortices, that were entered in the subsequent shape analysis, together with the three anatomical subregions of the striatum (caudate nucleus, putamen and nucleus accumbens; c.f. Fig 2b).

**Figure 2:**
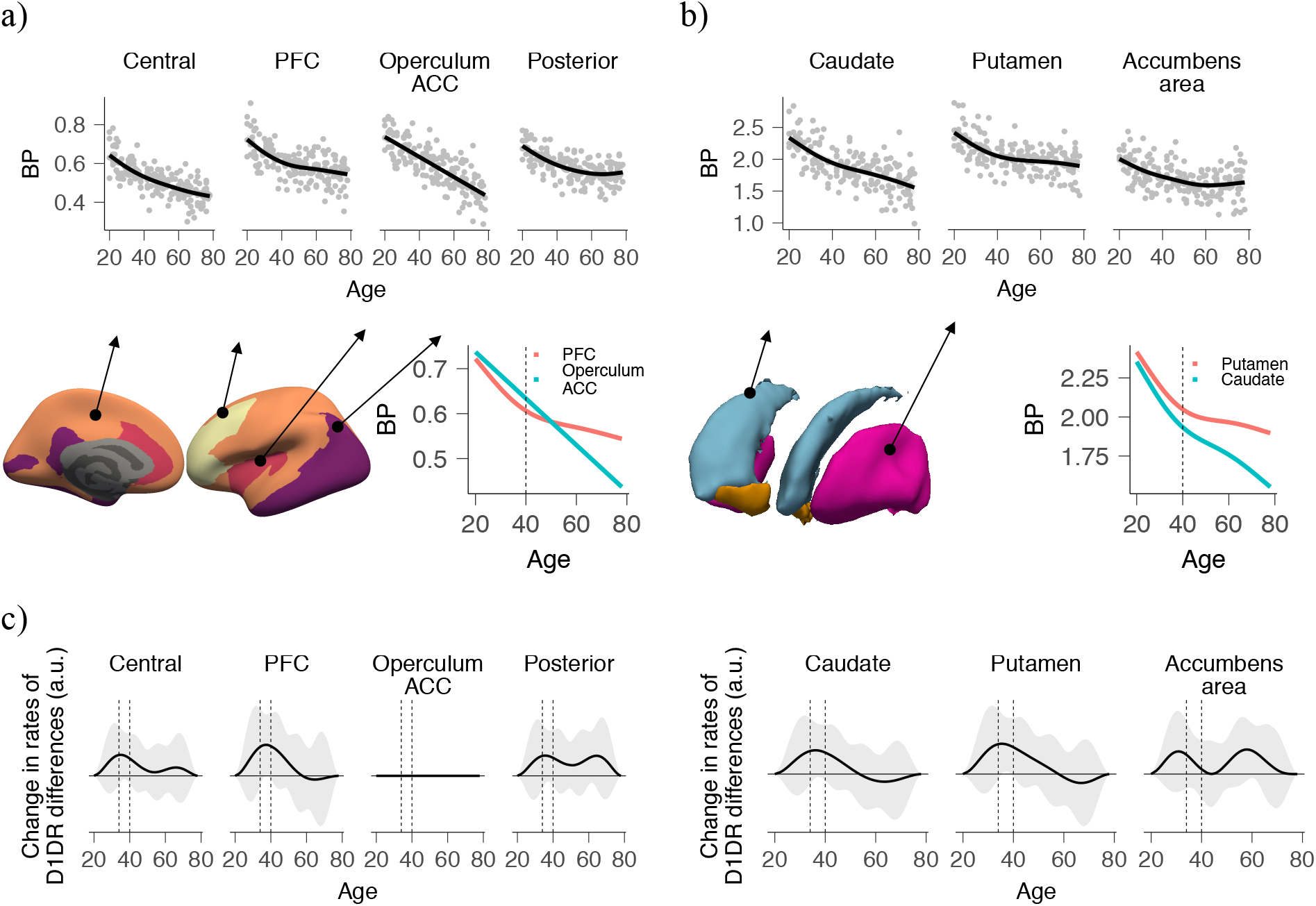
Age-related differences in [^11^C]SCH23390 BP_ND_ across the adult lifespan (age 20 – 80 years, n=176, 50% females, ∼15 / 15 M/F participants per decade). **A**: Cortical data. Scatter plots represent mean ROI-wise D1DR availability in relation to age, while the solid lines represent GAM fits. Data are shown in a reduced set of ROIs based on similar age-related trajectories within Desikan-Killiany parcellation. Lower panel depicts aggregated cortical ROIs, and an overlay of PFC- and cingulo-opercular-related age-trajectories highlighting a regionally deviant pattern from approximately 40 y (dashed line). **B**: Striatal data. Similar to the cortex, regionally distinctive patterns in age-trajectories of caudate and putamen were observed. **C**: Change in rates. D1DR differences were assessed using second-order GAM-derivatives. Putamen, PFC, and caudate indicated a peak at approximately age 35 – 40 y (dashed vertical lines), providing statistical evidence of non-monotonic age-trajectories in the putamen and PFC (c.f. 95% CI).

Model comparison using Akaike information criteria (AIC) (Schwarz, 1978) was used to identify the best age model fit. Log-transformed regional D1DR trajectories were best explained by a non-linear GAM in all ROIs, except caudate and operculum & ACC (Table 1, ΔAIC range = 2 - 13), indicating non-linear age-related effects in the putamen, prefrontal cortex, central and posterior cortices, and nucleus accumbens D1DR. Notably, log-transformation of the D1DR scores ensured that the non-linearities were not due to monotonous exponential decay – as suggested in some past studies (Jucaite et al., 2010; Suhara et al., 1991). Further analyses revealed an inflection point near age 40 y in several regions (Fig 2a-b). In accordance, a significant peak at approximately 35 – 40 years of age in the second-order GAM-derivatives (Pedersen et al., 2019) of putamen and PFC D1DR age trajectories was found (Fig 2c), corroborating a change in the rates of reduction close to this age period. In further support of a changing degree of age dependency before and after age 40, a bi-linear model incorporating different age-related slopes in early (age 20 – 40 y) versus later adulthood (age ≥ 40y) outperformed a constant linear model (and GAM) in several regions (Table 1).

**Table 1:** Akaike information criteria (AIC) scores of regional models of D1DR age trajectories. Log-transformed D1DR availability served as dependent variable to account for putative constant relative rates of decline (exponential decay model). Linear model with constant slope across the adult life span served as a reference model. General additive model (GAM) allowed for assumption-free analysis of nonlinear age effect. A bi-phasic linear model incorporating different slopes of age-related differences in early (age 20 – 40 y) versus later adulthood (age ≥ 40y) assumes a change in the slopes at a particular age (40 y). Lower AIC indicated superior model fit (improved data fit relative to a linear model are highlighted in bold font).

|  | AIC score |  |  |
| --- | --- | --- | --- |
|  | Linear | GAM | Bi-linear |
| Putamen | -289 | **-297** | **-299** |
| Caudate | -203 | -202 | **-204** |
| Accumbens | -198 | **-205** | **-202** |
| PFC | -247 | **-252** | **-253** |
| Operculum & ACC | **-217** | -216 | -215 |
| Central | -255 | **-257** | **-257** |
| Posterior | -305 | **-318** | **-316** |

Next, to quantify percent D1DR differences per decade in the two age segments – early adulthood (age < 40 y) and midlife and beyond (age ≥ 40 y) – bi-linear modeling with bootstrapping was conducted. In accord with non-linear age trajectories, the relative rates of D1DR reduction were higher in early adulthood as compared to midlife and late adulthood (Fig 3) – with the exception of operculum and ACC which showed a linear relation with age. Furthermore, in early adulthood the cross-regional estimates of % D1DR difference per decade were within 3.0 %-units and all 95% confidence intervals (CI) were overlapping, whereas several regions were significantly different from one another in midlife and beyond (Fig 3, non-overlapping 95% CIs). This regional heterogeneity post midlife was further characterized by two sets of regions exhibiting dissimilar rates of difference. The first included operculum and ACC, caudate, and central cortex, characterized by larger relative D1DR differences (mean [95% CI] = -6.2 % [-7.7, -4.6]) compared to the second set of regions including putamen, PFC, posterior cortex and nucleus accumbens (mean [95% CI] = - 1.8 % [-3.5, 0.1]).

**Figure 3:**
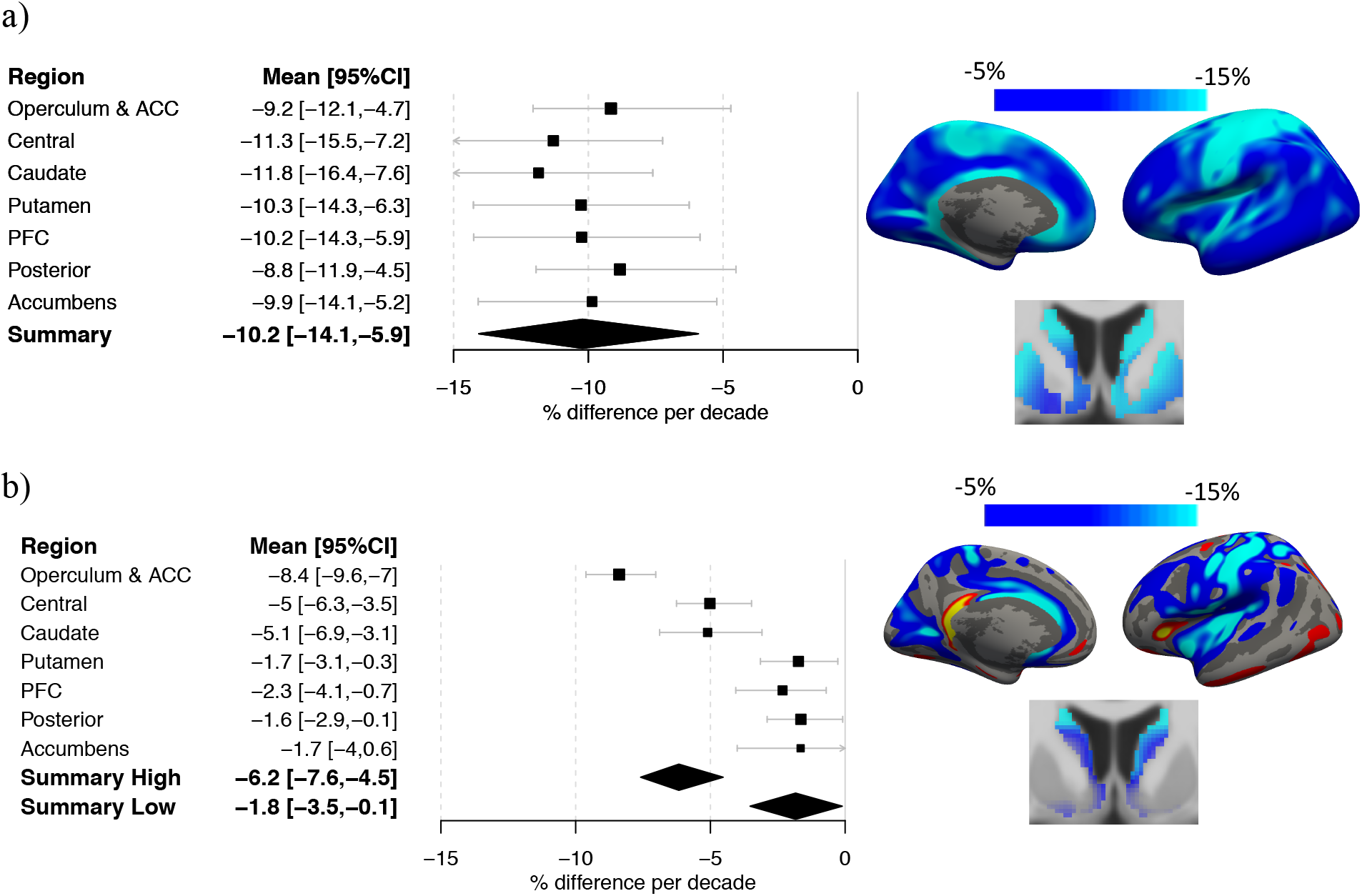
Bi-linear analysis of age-related D1DR differences. ROI- (left) and voxel-based (right) percent D1DR difference per decade are reported. **A**: results in early adulthood (age 20 – 40 y). **B**: results in later adulthood (age ≥ 40 y). D1DR % difference per decade was calculated as %DPD = 100% (BP_x_/BP_y_ - 1)/#decades, where BP_x/y_ are predicted estimates of BP_ND_ at [y=20, x=40] and [y=40, x=80], and #decades was 2 and 4, respectively. Bootstrapping (n=500) was used for estimation of mean (box) and 95% CI (whiskers) of %DPD. Non-overlapping CIs were interpreted as significant rate differences (p < 0.05).

### Associations of D1DR and brain health in striatal subregions

We next investigated the relation between white matter lesion volumes and interregional differences in the effects of aging. We hypothesized that cerebrovascular burden – as indexed by white matter lesion (WML) volume (Schmidt et al., 2012) – may be more closely coupled with caudate than putamen D1DR availability (Karalija et al., 2019), and modulate late life differences in age-related trajectories for DA-rich regions (c.f. Fig 2 & 3). WML volume ranged between 0 - 27.0 mL for the sample, with negligible WML manifestation in early adulthood (volume range 0 – 1.2 mL, n = 60). In late adulthood (age ≥ 40 y, n = 116) WML volume was more strongly coupled with caudate than putamen D1DRs (Fig 4, age-partial correlations: *r*_*caudate*_ = -0.29, p < 0.01; *r*_*putamen*_ = -0.15, n.s., test for difference in dependent correlations *t* = -2.24, p = 0.03). Notably, inclusion of a region-specific effect of WML volume in a hierarchical model of D1DR differences moderated interregional differences in age slopes in the middle age and beyond (Supplementary Table 5), suggesting that the interregional differences in the age trajectories were related to distinct cerebrovascular influences to D1DR availability in striatal subregions. Predicted age-related trajectories of caudate and putamen D1DR availability (Fig 4b) demonstrated a high degree of similarity between the striatal subregions after adjustment for WML volume.

**Figure 4:**
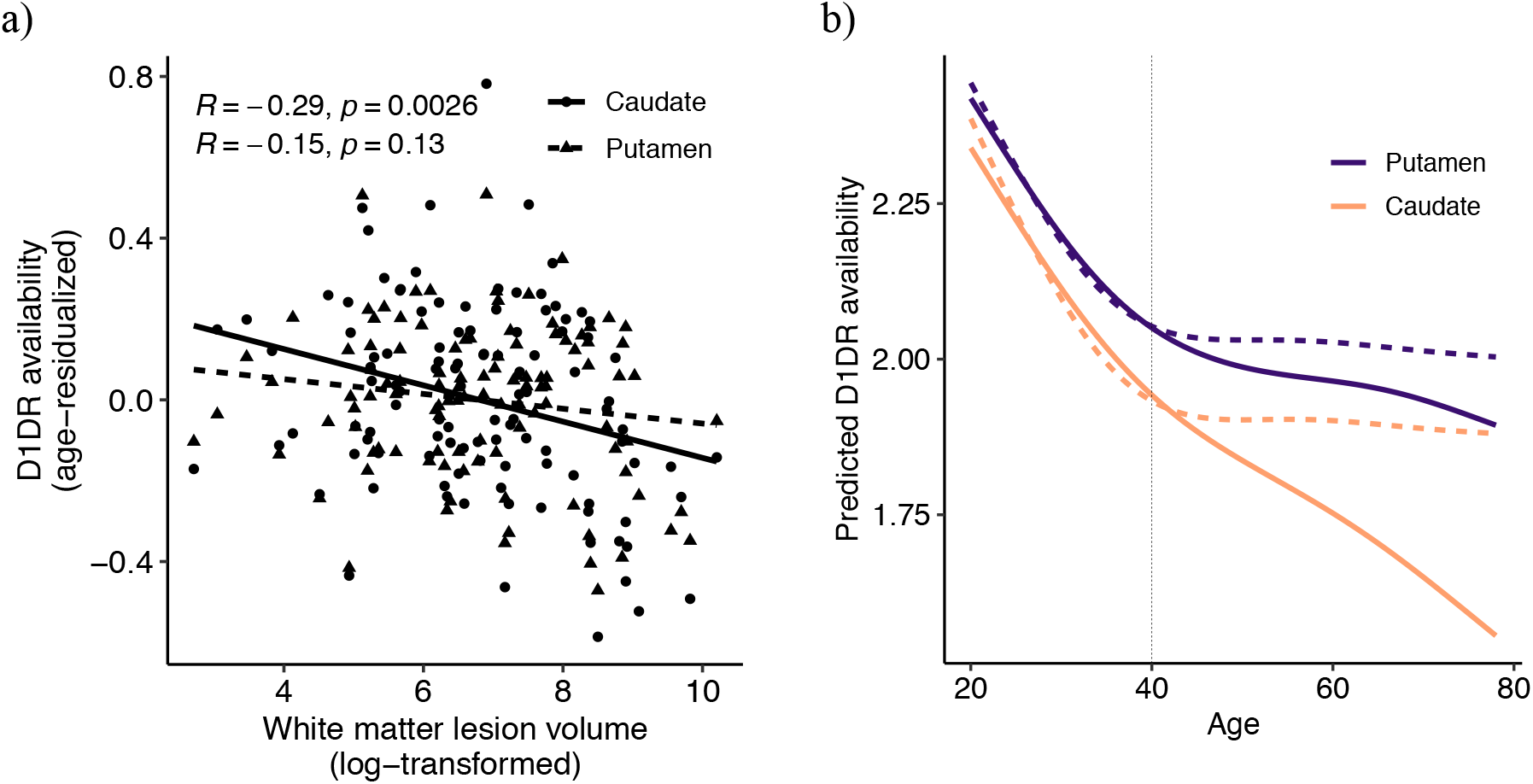
Regionally distinctive relations between D1DR availability and cerebrovascular burden (white matter lesion volume). **A**: scatter plot showing a stronger partial correlation between caudate than putamen D1DR and WML volume. **B**: Predicted age trajectories of putamen and caudate D1DR availability. Interregional differences in age-related trajectories of raw data (solid lines) were abolished after adjustment for the region-specific effect of white matter lesion volume (dashed lines).

### Associations of caudate D1DR and neurocognitive function across the adult lifespan

We next investigated the implications of caudate D1DR availability on cognition. A composite score of episodic and working memory was used (see Materials & Methods), given that caudate DA receptors have previously been shown to be related to memory processing (Bäckman et al., 2011; Nyberg et al., 2016). We hypothesized that high D1DR levels in early adulthood (age < 40y) – if reflective of protracted D1DR development – may relate to excessive DA modulation, low strength of local caudate-to-cortex functional connectivity (FC; see Mat. & Met.), and have negative implications for memory. In late adulthood (age ≥ 40y), in contrast, the low D1DR levels may relate to insufficient DA modulation, elevated FC levels, and have negative effects on memory.

Lower caudate D1DRs was associated with less efficient memory function in older (Pearson’s *r*_*old*_ = 0.44, *p* < 10^−6^; age-partial *r*_*old*_=0.24, *p* < 0.01), but there was no significant association in younger adults (Pearson’s *r*_*young*_ = 0.15, *p* = 0.28; age-partial *r*_*young*_=0.15, *p* = 0.26). Although the relation between caudate D1DR and memory was only moderately modulated by age (Supplemental Fig 3), this finding suggests that the relation between D1DR and memory in young adulthood may differ from that observed in older age. Similar to D1DRs, greater caudate-to-cortex FC was positively linked to memory in younger (Pearson’s *r*_*young*_ = 0.22, *p* = 0.10; age-partial *r*_*young*_=0.22, *p* = 0.10), but negatively in older adults (*r*_*old*_ = - 0.32, *p* < 0.001; age-partial *r*_*old*_ = -0.19, *p* = 0.04; Supplemental Fig 3, Fisher’s z-test for difference in correlations p < 0.05). This finding suggests that excessively low (in early adulthood) and excessively high (in late adulthood) caudate FC were suboptimal in terms of memory function.

We then examined whether D1DR in different age-segments and at different levels of FC exhibited distinct relations with memory. Analysis of variance (ANOVA) stratified by age-segment (age < 40y, age ≥ 40y), revealed a negative effect of high D1DR availability on memory in early adulthood when concomitant with low FC (Figure 5a), a finding that was in stark contrast to the other strata, where higher D1DR levels generally predicted better memory (two-way ANOVA: D1DR x FC, *F* = 5.6, *p* < 0.05, age < 40y). The association between D1DR level and memory was thus inversed as a function of FC level in young adults (c.f. Fig 5a), in support of the hypothesis that high D1DR may be associated with suboptimal memory function when coupled with the manifestation of excessive DA modulation (i.e. attenuated FC). In contrast, in older age, low D1DR level coupled with the manifestation of insufficient DA modulation (i.e. elevated FC) was associated with low memory scores. Collectively the present findings supports an inverted-U shaped pattern in the effects of D1DR on memory across the adult life span (Fig 5b).

**Figure 5:**
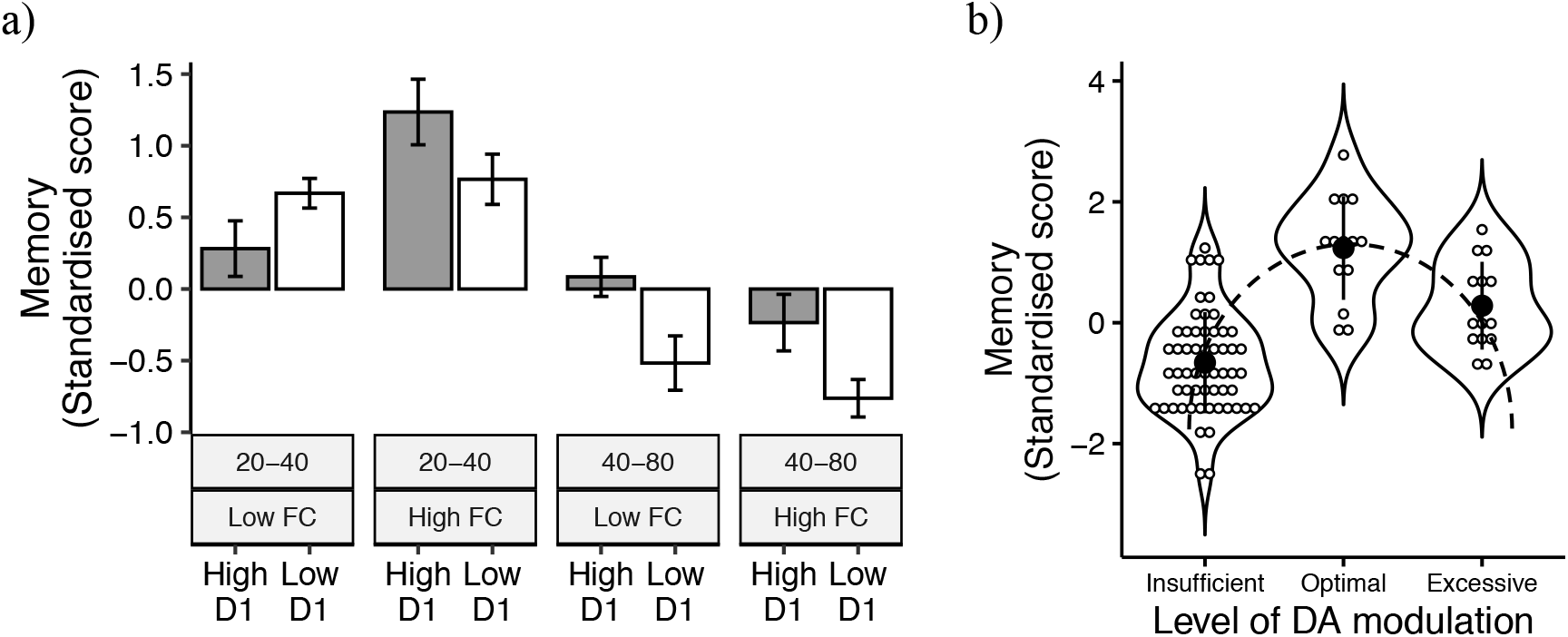
Analysis of memory scores in relation to caudate D1DR availability and caudate-to-cortex functional connectivity (FC). **A**: analysis of memory scores (mean ± se) across age-segments (age < 40y, age ≥ 40 y), and across D1DR x FC groups (median-split; high/low). Disparate association to memory was observed in young individuals with high D1DR but low FC as compared to other groups. In early adulthood high D1DR level in combination with low FC was associated with lower memory scores, whereas in other groups high D1DR tended to associate with high memory scores [two-way (D1DR x FC level) ANOVA p<0.05, early adulthood]. **B**: memory scores in relation to the postulated level of DA modulation followed an inverted-U shape pattern (dashed line illustrates the pattern). The DA modulation levels were labelled by: Excessive = age < 40, high D1, low FC; Optimal = age < 40, high D1, high FC; Insufficient = age ≥ 40, low D1.

## Discussion

The present study provides *in vivo* evidence of a bi-phasic pattern of D1DR differences across the adult lifespan. A *post mortem* study found a bi-phasic D1DR age-pattern during childhood (Seeman et al., 1987), but it has remained elusive whether age effects in early adulthood (Bäckman et al., 2011; Montague et al., 1999; Rinne et al., 1990; Seeman et al., 1987; Suhara et al., 1991) have a different origin compared to midlife and later age segments. An inflection point in D1DR age trajectories provides novel evidence that a transition between periods takes place during adulthood. Importantly, the age-related D1DR differences were generally much larger in early than in later age segments. Furthermore, the age-related differences were characterized by regional homogeneity in early but regional heterogeneity in late adulthood. These findings resonate well with cross-regional synchrony of developmental events (Bakken et al., 2016; Lidow et al., 1991), and regionally specific susceptibility to degenerative events in later life (e.g. (Iadecola, 2013)), respectively. A multifactor model of D1DR alterations across the adult lifespan like that proposed for brain morphology (Fjell et al., 2013; Sowell et al., 2003) and connectivity (Bartzokis et al., 2012; Wang et al., 2012), provides a biologically plausible account of the present findings.

A tentative multifactor model posits D1DR reductions in early adulthood to reflect protracted receptor development, followed by selective receptor degeneration in older age. The alternative – and currently prevailing – unimodal account assumes a continuum of negative age-related consequences with early onset in young adulthood. Speaking against the unimodal account, the age-related trajectories in a number of regions – including putamen and prefrontal cortex – decelerated markedly in midlife. This is in apparent discord with typically accelerating – not decelerating – rates of age-related decline for cognition and brain integrity (Fjell et al., 2013; Raz et al., 2005; Rönnlund et al., 2005; Schaie, 1994). In support of protracted receptor development, extant evidence from samples including juveniles indicate an extraordinarily late peak in D1DR (over)expression during the third decade of life (Rothmond et al., 2012; Weickert et al., 2007). Together with findings of declining levels of D1DR markers in adolescence and beyond (Jucaite et al., 2010; Seeman et al., 1987), it is likely that contributions of protracted D1DR development extend to early adulthood.

In keeping with adult D1DR development, a negative relation between high D1DR expression and memory was found in the youngest participants. This finding conforms with the inverted-U shaped model of lifespan age differences in DA modulation (e.g. (Li et al., 2010)). By this account, overexpression of D1DR during development is thought to contribute to excessive DA modulation – as indicated here by low striatal functional connectivity – and poorer cognition. Notably, the peak of excessive DA modulation has been assumed in adolescence (e.g. (Li et al., 2010)), but here, evidence in support of the inverted-U shaped pattern was found in an adult sample. Thus, the present findings suggest that functionally relevant DA development may extend further into early adulthood than previously reported. In line with our findings, a recent longitudinal study reported developmental change in D2DR availability in early adulthood (18 – 32 y) (Larsen et al., 2020). Others have also found more drastic D2DR and DAT reductions in early compared to late adulthood (Antonini and Leenders, 1993; Mozley et al., 1999; Yeh et al., 2022), collectively pointing to the possibility of ongoing development of the DA system in adult age.

Not all regions followed a bi-phasic age pattern. Most notably, caudate and cingulo-opercular regions exhibited linear age-related trajectories. Nevertheless, analysis of the rates of age-related D1DR differences showed no apparent cross-regional divergence in early adulthood, but rather, the divergence was manifested in the later age segments. This dichotomy may indicate the presence – and predominance – of a common cause (i.e. development) during early adulthood which wanes by middle age. From middle age and beyond the predominance is shifted towards negative age-related consequences, with likely regionally selective features (e.g. (Iadecola, 2013; Seaman et al., 2018)). By this account, the bi-phasic age pattern could hence have been moderated by the onset of negative age-related consequences overlapping with the end of receptor development. In keeping with this view, selective associations with midlife cerebrovascular burden were observed in the striatal subregions. Notably, the age-related trajectory of caudate D1DR was indistinguishable from that of putamen when adjusting for the cerebrovascular burden, speaking for a bi-phasic age-trajectory also in the caudate when adjusted for vascular factors. The cingulo-opercular region co-localize with past observations of exacerbated reductions in DAT (e.g. Kaasinen et al., 2015), brain glucose metabolism (van Aalst et al., 2021), and cortical thickness (Elliott et al., 2021). Hence, it seems plausible that the midlife D1DR reductions are linked to alterations of other brain markers within these regions. Unlike caudate, no region-specific predictors of cortical D1DR reduction were however identified. Therefore, it remains unknown whether the cingulo-opercular regions would also show a bi-phasic pattern if adjusting for the (unidentified) co-variates of brain decline.

Regardless of the precise nature of these midlife D1DR losses, the regions involved are implicated in cognition. The cingulo-opercular areas, central components of the salience network (Dosenbach et al., 2008), have been linked to sustained attention (Sadaghiani and D’Esposito, 2015) and allocation of executive control (Shenhav et al., 2013). Caudate, in concert with cingulo-opercular regions and midbrain, has been implicated in learning and biasing value-based action policies (Jiang et al., 2015; Westbrook and Braver, 2016). Toward this end, a coupling between D1DR availability in dorsal striatum and action learning has been found (de Boer et al., 2019). These functions in turn tend to rely on signals of reward (Yee et al., 2021; Yee and Braver, 2018), typically attributed to the mesocorticolimbic DA pathway (Schultz, 1998). Therefore, it is conceivable that the age-related D1DR reductions in these regions might bear specific importance to motivation-based modulation of cognition (De Boer et al., 2017; Westbrook and Braver, 2016; Yee et al., 2021; Yee and Braver, 2018), beyond concomitant changes in structure and metabolism. The present findings are hence in keeping with neurocognitive theories of aging that attribute losses in cognitive control to deficient DA modulation (Braver and Barch, 2002).

Our findings of moderate rate of D1DR differences in midlife and beyond stand in stark contrast to those estimated in past imaging studies (for a review see Karrer et al. (2017)), but are in a good agreement with post-mortem investigations (mean = -4.0 % per decade) (Rinne et al., 1990; Seeman et al., 1987). Notably, the present analysis departs from the majority of past research by using a non-monotonous age model, in contrast to the typically utilized polynomial or monotonous exponential decay models. The predominance of monotonous age models in prior work likely reflects the paucity of well-powered investigations, but also methodological limitations in DA assessment. In contrast to the rigorous control of partial-volume-effects (PVE) in the present study, the majority of past research has likely been affected by age-related differences in concomitant brain atrophy. In particular, concomitant brain atrophy exacerbates suppression of the PET signal, inducing negative biases in the rate of age-related alterations of DA markers (Greve et al., 2016; Smith et al., 2019). As a result, the current data may match better with extant *post mortem* than *in vivo* investigations. A further concern is related to the health of elderly participants in past research. Whereas the present study carefully randomized participants and screened for brain health (Nordin et al., 2022), this was not necessarily carried out in all past research. Indeed, some prior studies have included healthy participants scanned for diagnostic purposes (e.g. Kaasinen et al., 2015) and, despite lack of clinically relevant findings for the particular DA marker, it is possible that these participants may not reflect typical healthy aging. In line with this reasoning, some investigations found no age effect for DA in later adulthood (e.g. (Jakobson Mo et al., 2013; Reeves et al., 2005)), or bi-linear age-related patterns across early and late adulthood (Antonini and Leenders, 1993; Mozley et al., 1999; Yeh et al., 2022). That said, the elderly participants in the present study were not “super-agers”, as indicated by typical group-level patterns of age-related cognitive deficits (Nordin et al., 2022). Taken together, present findings suggest that brain-wide DA losses in healthy aging may be substantially smaller than previously reported.

The cross-sectional design of the present study warrants certain cautions. The type of analysis presented here does not pertain to individual age-trajectories but assumes that the groups across different age-segments present a good reflection of what may have happened at the individual level. Efforts were made in recruitment to avoid disbalanced cohorts, and rigorous post-screening found no evidence for unexpected or dramatic differences across the age- or sex-groups (Nordin et al., 2022). Nevertheless, longitudinal follow-up studies are warranted to confirm that between-person associations reflect within-person trajectories.

## Conclusions

The present findings provide novel insights into distinguishable patterns of D1DR age-differences across the adult lifespan that putatively reflect disparate developmental events and effects of ageing. The findings can inform future studies of neurocognitive development in health and disease, as it is currently not well understood how atypical DA age-trajectories differ from typical development. Furthermore, this work may inform studies of how interindividual DA differences relate to cognition across the adult lifespan. In particular, evidence of excessive DA modulation in adulthood – in contrast to adolescence exclusively – illustrates an equivocal, inverted-U shaped pattern between DA function and cognition across the adult lifespan.

## Methods

We have reported the DyNAMIC study design, recruitment procedure, imaging protocols, cognitive testing and lifestyle questionnaires in detail elsewhere (Nordin et al., 2022). DyNAMIC is a prospective study of healthy individuals across the adult lifespan. Here, we restrict the description to the methodological and material aspects of direct relevance to the present study. The study was approved by the committees of Ethical and Radiation Safety, and all participants gave written informed consent prior to testing.

### Participants

The sample was based on randomly selected invitees from the population registry of Umeå, Sweden. Responders were screened for exclusion criteria, including contraindications to magnetic imaging, mini mental state examination (MMSE) ≤ 25, psychoactive medication, history of mental illness, and brain abnormalities. One hundred eighty (n=180) volunteers were included in the study in an age- and sex-balanced manner across 20 – 80 y (n=30 per decade, 50% female). 177 participants completed PET scanning using [^11^C]SCH23390, but in one participant there were indications of subcutaneous injection leading to exclusion of this participant; other reasons for drop-out from PET were technical problems, and one participant declined to participate in PET.

### Imaging Procedures

Magnetic resonance (MR) imaging was conducted using a 3 tesla scanner (Discovery MR 750, General Electric), and a 32-channel phased-array head coil. PET scanning was conducted using a hybrid PET/CT system (Discovery PET/CT 690, General Electric).

### PET Imaging

Production of [^11^C]SCH23390 was performed by the radiochemistry laboratory of Norrlands Universitestsjukhus, Umeå University. Target radioactivity in intravenous injections of [^11^C]SCH23390 was 350 MBq (range = [205, 391] MBq, mean ± sd = 337 ± 27 MBq, no significant differences across age-groups (age < 40 and age ≥ 40 y), Student’s t-test *t* = 0.79, n.s.). Participants were positioned on the scanner bed in supine position, and individually fitted thermoplastic masks were used to prevent excessive head movement. Preceding the injection, a 5-min low-dose helical CT scan (20 mA, 120 kV, 0.8 s per revolution) was obtained for PET-attenuation correction. Continuous PET-measurement in list-mode format was initiated at the time of injection and continued for 60 minutes. Offline re-binning of list-mode data was conducted to achieve a sequence of time-framed data with increasing length: 6 × 10, 6 × 20, 6 × 40, 9 × 60, 22 × 120 s (a total of 49 frames). Time-framed, attenuation-, scatter-, and decay-corrected PET images (47 slices, 25 cm field of view, 256×256-pixel transaxial images, voxel size 0.977×0.977×3.27 mm3) were reconstructed using the manufacturer-supplied iterative VUE Point HD-SharpIR algorithm (6 iterations, 24 subsets, resolution-recovery).

Estimation of target binding potential (BP) relative to non-displaceable (BP_ND_) binding as measured in the cerebellum was conducted according to procedures described previously (Nordin et al., 2022). Frame-to-frame head motion correction (translations range = [0.23, 4.22] mm, mean ± sd = 0.95 ± 0.54 mm, a trend-level difference across age-groups (age < 40 and age ≥ 40 y), Student’s t-test *t* = 2.0, p = 0.047, mean ± sd young = 1.07 ± 0.52, mean ± sd old = 0.90 ± 0.55), and registration to T1-weighted MRIs were conducted by using Statistical Parametric Mapping software (SPM12, Wellcome Institute, London, UK), and corrected PET data were re-sliced to match the spatial dimensions of MR data (1 mm^3^ iso-tropic, 256 × 256 × 256). Partial-volume-effect (PVE) correction was achieved using the symmetric geometric transfer matrix (SGTM; regional correction) or Muller-Gartner (voxel-wise correction) method implemented in FreeSurfer (Greve et al., 2016), and an estimated point-spread-function of 2.5 mm full-width-at-half-maximum (FWHM). Regional estimates of BP_ND_ were calculated within Desikan-Killiany cortical parcellation (Desikan et al., 2006) and subcortical regions as provided in automated FreeSurfer segmentations (41 regions-of-interest), using the simplified reference tissue model (SRTM; (Lammertsma and Hume, 1996)). Voxel-wise estimates of BP_ND_ were calculated within gray-matter voxels (GM probability > 0.9) using the multi-linear (simplified) reference tissue model (MRTM), with fixed k_2_’ (MRTM2; (Ichise et al., 2003)). Voxel-wise BP_ND_ maps were spatially normalized to match Montreal Neurological Institute (MNI) space using DARTEL-derived deformation fields. Cortical surface-reconstructions of BP_ND_ maps in MNI-space were conducted using FreeSurfer.

### MR Imaging

High-resolution anatomical T1-weighted images were collected using a 3D fast spoiled gradient-echo sequence. Imaging parameters were as follows: 176 sagittal slices, thickness = 1 mm, repetition time (TR) = 8.2 ms, echo-time (TE) = 3.2 ms, flip angle = 12º, and field of view (FOV) = 250 × 250 mm. Anatomical T1-weighted images were used to parcel cortical and subcortical structures with the Freesurfer 6.0 software (https://surfer.nmr.mgh.harvard.edu; (Fischl et al., 2002)). Striatal volumes were manually corrected using the Voxel Edit mode in Freeview when necessary.

Whole-brain functional images were acquired during naturalistic viewing (movie watching). Functional images were sampled using a T2*-weighted single-shot echo-planar imaging (EPI) sequence, with a total of 350 volumes collected over 12 minutes. The functional sequence was sampled with 37 transaxial slices, slice thickness = 3.4 mm, 0.5 mm spacing, TR = 2000 ms, TE = 30 ms, flip angle = 80º, and FOV = 250 × 250 mm.

fMRI preprocessing was carried out using the Statistical Parametric Mapping software package (SPM12; Wellcome Department of Imaging Science, Functional Imaging Laboratory). The functional images were first corrected for slice-timing differences, motion, and signal distortions and the time series were subsequently demeaned and detrended followed by simultaneous nuisance regression and temporal high-pass filtering (threshold at 0.009 Hz) in order not to re-introduce nuisance signals (Hallquist et al., 2013). Nuisance regressors included average CSF and WM time-series and their derivatives, 24-motion parameters (Friston et al., 1996), a binary vector flagging motion-contaminated volumes exceeding framewise displacement (FD) of 0.2 mm (Power et al., 2012), in addition to an 18-parameter RETRICOR model (Glover et al., 2000; Hutton et al., 2011) of cardiac pulsation (up to third-order harmonics), respiration (up to fourth-order harmonics), and first-order cardio-respiratory interactions estimated using the PhysIO Toolbox v.5.0 (Kasper et al., 2017). Nuisance-regressed images were subsequently normalized to a sample-specific group template (DARTEL; (Ashburner, 2007)), spatially smoothed using a 6-mm FWHM Gaussian kernel and affine-transformed to stereotactic MNI space.

Functional connectivity (FC) graphs were created by correlating average time series sampled from 264 cortical locations ((Power et al., 2011); 5-mm spheres) in addition to two putative ROIs in the right and left caudate (MNI coordinates: right [10, 4, 2], left [-12, 12, 6]). Following Fisher’s r-to-z transformation, a bilateral estimate of caudate FC strength was computed for each subject as the average sum of positive edge weights for the two vertices. For assessment of white-matter hyperintensities, a fluid-attenuated inversion recovery (FLAIR) sequence was acquired. A total of 48 slices were obtained with a slice thickness of 3 mm, TE = 120 ms, TR = 8000 ms, TI = 2250 ms, and FOV = 240 × 240 mm. The lesion segmentation tool (LST; https://www.applied-statistics.de/lst.html) in SPM12 was used to segment white-matter lesions (hyperintensities in FLAIR).

### Memory

Tests of episodic memory included word recall, number-word recall, and object-location recall. In word recall, participants were presented with 16 words that appeared one by one on the computer screen. During the first phase, participants encoded each word for 6 s, and following presentation of all items in the series, participants used the keyboard to type in as many of the words they could recall, in any order. Performance was defined as the number of correctly recalled words. In number-word recall, participants were required to memorize pairs of 2-digit numbers and concrete plural nouns (e.g. 46 dogs). Ten number-word pairs were presented consecutively, each displayed for 6 s. Retrieval immediately followed, in which every word was consecutively presented again, but in a different order than during encoding. For each word, participants had to recall the associated 2-digit number, and type the answer using the keyboard. A response was required for each word, meaning that participants had to provide a guessing-based response even if they did not recall the correct number. For object-location memory, participants encoded objects presented on different locations in a 6 × 6 square grid displayed on the computer screen. Each encoding trial involved 12 objects, presented one by one, in distinct locations within the grid. Each object-position pairing was displayed for 8 s before disappearing. Directly following encoding, all objects were simultaneously displayed next to the grid for participants to move them (in any order) to their correct location in the grid. If unable to recall an object’s correct position, participants had to guess and place the object at a location to the best of their ability. A composite score of episodic memory (EPM) was computed. First, summary scores per task were computed across blocks or trials of tasks. Next, these summary scores were standardized (T-score: Mean = 50; SD = 10), and finally, a composite score was created by averaging the T-scored measures of each task.

Working memory was tested using three tasks, letter updating, number updating, and spatial updating. During letter updating, participants were presented with a sequence of capital letters (A– D), consecutively on the computer screen, requiring them to update and to keep the three lastly presented letters in memory. The letters were presented for 1 s, with an ISI of 0.5 s. When prompted, which could be at any given moment, participants provided their response by typing in three letters using the keyboard. Four practice trials were completed by all participants, followed by 16 test trials consisting of either 7-, 9-, 11-, or 13-letter sequences. Across all 16 trials, the maximum number of correct answers were 48 (16 trials × 3 reported letters = 48). The number-updating task had a columnized numerical 3-back design. Three boxes were present on the screen throughout the task, in which a single digit (1– 9) was presented one at a time, from left to right during 1.5 s with an ISI of 0.5 s. During this ongoing sequence, participants had to judge whether the number currently presented in a specific box matched the last number presented in the same box (appearing three numbers before). For each presented number they responded yes/no by pressing one of two assigned keys (“yes” = right index finger; “no” = left index finger). Four test trials, each consisting of 30 numbers, followed after two practice trials. Performance was defined as the sum of correct responses across the four test trials, after discarding responses to the first three numbers in every trial (as these were not preceded by any numbers to be matched with). The maximum score was 108 (27 numbers × 4 trials). In the spatial-updating task, three 3 × 3 square grids were presented next to each other on the computer screen. At the beginning of each trial, a blue circular object was, at the beginning of each trial, displayed at a random location within each grid. Following a presentation time of 4 s, the circular objects disap-peared, leaving the grids empty. An arrow then appeared below each grid, indicating that the circular object in the corresponding grid was to be mentally moved one step in the direction of the arrow. The arrows appeared stepwise from the leftmost grid to the rightmost grid, each presented for 2.5 s (ISI = 0.5 s). The exercise of mentally moving the circular object was repeated one more time for each grid, prompted by three new arrows, resulting in the object having moved two steps from its original location at the end of each trial. Using the computer mouse, participants then indicated which square the circular object in each grid had ended up in. If unsure, they provided guesses. The test was performed across 10 test trials, preceded by five practice trials. Performance was calculated as the sum of correct location indications across trials, with a maximum score of 30. A composite score of working memory (WM) was computed based on the three test scores, in the same way as described for EPM.

Finally, a composite score of memory function was calculated using principal component analysis (PCA) of WM and EPM scores across all participants (n = 176). Unrotated PCA loadings of the first component were 0.9 for both WM and EPM, and the first PC scores were extracted to indicate memory function.

### Statistical Analyses

Generalized additive modeling (GAM; (Wood, 2017)) allowing for smooth functions was used as the primary age model, and model comparisons using Akaike information criteria (AIC; (Schwarz, 1978)) were conducted to decide which model had most empirical support. Past research has predominantly considered linear and exponential decay D1DR – age models (Karrer et al., 2017; Suhara et al., 1991), the latter reflecting a non-linear, but monotonous, function of constant decay in relation to the concentration, whereas the preceding model is a good approximation of such decay when the decay rate (λ) is very low. Here, we hypothesized non-monotonous non-linear relationships between age and D1DR, which are poorly characterized by linear or exponential decay models. Conformity to the exponential-decay model was assessed using log-transformed D1DR availability and a linear model.

All ROI-level analysis were conducted using R (version 4.0.3). Univariate outliers in regional D1DR availability were searched and removed within each decade and sex (n ≈ 15), by using the box-plot method (version 4.0.3). Outlier BP_ND_ were generally related to poor PET-model fit, and bilateral exclusion of BP_ND_ was present in less than 1 % of the aggregate ROIs. R-package mgcv (version 1.8-33) was used for GAM analysis, and smoother functions in D1DR – age models were configured with six (k = 6) initial thin-plate (tp) basis functions, to avoid overfitting (method = REML, family = Gaussian). Complexity of the smoothers were penalized (Wood, 2017), and the resulting effective degrees of freedom were allowed to vary across regions. R-package gratia (version 0.6.0) was used for computation of GAM derivatives (1^st^ and 2^nd^ order) and the corresponding confidence intervals (CI 95%). Hierarchical (multivariate) GAMs (Pedersen et al., 2019) were constructed for the purpose of 1) dimension reduction in cortical ROIs; and 2) testing interregional differences in absence and presence of covariates. For the first, a HGAM including cortical D1DR availability (34 ROIs) as the dependent variable, and a global (fixed) effect of smooth age, regional (random) effects of intercept and smooth age, and a random effect of individual (intercept) as predictors was constructed. In this model, the global effect of age from cortical age trajectories was regressed out, and residual regional age effects were piped into a shape-respecting K-means clustering algorithm (R-package kmlShape, version 0.9.5; (Genolini et al., 2016)), to find regions sharing similar trajectories. Number of clusters were empirically optimized at k = 3. For the second, HGAMs were configured across regions (pairs), where the preceding analysis had suggested regionally-variant age trajectories; using a likelihood-ratio test, a model with a regional (random) effect of smooth age beyond a common smooth effect of age was compared to the common age-effect model. A significant result (p<0.05) in the likelihood-ratio test indicated regional differences in age trajectories (Pedersen et al., 2019). Furthermore, regional (random) effects of additional predictors (WML, GM, WM volumes) were used to test the persistence of regionally-variant age effects in the presence of regionally-variant exogenous predictors. Contributions of additional variables to regional D1DR differences beyond the effects of age were investigated using step-wise linear model comparisons assessing improvement in the model if the candidate variable was included (likelihood-ratio test, p<0.05). Shapiro-Wilk normality tests (skewness, kurtosis) were conducted, and log-transformed variables were used to ensure normally distributed variables when needed.

Percent D1DR difference per decade was used as a measure of reduction rate:

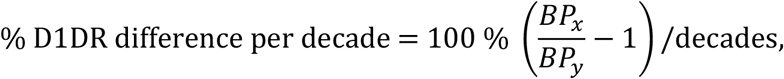

where *BP*_*x*_ is the BP_ND_ predicted at age *x*. R-package boot (version 1.3-25) was used for bootstrap-based estimation of the confidence interval % for D1DR differences per decade (n = 500 repetitions, CI 95% was used (Seaman et al., 2019)).

Voxel-wise PET analyses were conducted using Matlab (version 2017, Mathworks, US). Bi-linear age-models were fit using a piecewise-linear least-squares algorithm (https://www.mathworks.com/matlabcentral/fileexchange/40913-piecewise-linear-least-square-fit). Prior to model fit, voxel-wise maps of D1DR availability were smoothed using an edge-preserving Gaussian smoothing kernel (FWHM = 8 mm). Voxel-wise analyses were conducted in the entire sample, and for male and female participants.

Analysis of variance were conducted to investigate the interrelations between caudate-D1DR availability, caudate-FC, and memory scores. Two-way interaction effects were included, always controlling for the intercept effect of each variable.

