## Supplemental material for "Bi-phasic patterns of age-related differences in dopamine D1 receptors across the adult lifespan"

### ROI-based GAM-analysis of age-effect.

**Supplemental Table 1:** Summary of the univariate ROI-based GAM-analysis for the effect of (smooth) age. Percentage variance explained ($r^2$) and stars representing the Bonferroni-corrected p-values (*** p<0.001) of the age-term are reported. Percent BP difference was calculated as 100 x ($BP_{80}/BP_{20} – 1$), where 20 and 80 refer to the model-predicted $BP_{ND}$ at these ages (% BP difference per decade in parenthesis).

| | | $r^2$-adj. (smooth age) | | % BP difference (20 - 80 y) | |
|---|---|---|---|---|---|
| | **Region** | **Left** | **Right** | **Left** | **Right** |
| Subcortex | Caudate | 0.53**** | 0.40**** | -40.66 (-6.78) | -30.64 (-5.11) |
| | Putamen | 0.32**** | 0.34**** | -21.91 (-3.65) | -23.53 (-3.92) |
| | Accumbens | 0.25**** | 0.16*** | -20.41 (-3.40) | -16.66 (-2.78) |
| | Pallidum | 0.20**** | 0.03ns | 11.91 (1.98) | 14.59 (2.43) |
| | Hippocampus | 0.02ns | 0.01ns | -11.95 (-1.99) | -9.48 (-1.58) |
| | Amygdala | 0.06ns | 0.03ns | -13.70 (-2.28) | -0.21 (-0.04) |
| | Thalamus | 0.16*** | 0.16*** | 15.48 (2.58) | 26.45 (4.41) |
| Frontal | Caudal Middle Frontal | 0.27**** | 0.23**** | -26.94 (-4.49) | -30.69 (-5.11) |
| | Frontal Pole | 0.23**** | 0.21**** | -25.36 (-4.23) | -22.06 (-3.68) |
| | Pars Orbitalis | 0.21**** | 0.23**** | -24.87 (-4.15) | -24.75 (-4.13) |
| | Pars Triangularis | 0.29**** | 0.26**** | -25.94 (-4.32) | -23.15 (-3.86) |
| | Rostral Middle Frontal | 0.29**** | 0.29**** | -20.30 (-3.38) | -22.71 (-3.78) |
| | Lateral Orbitofrontal | 0.41**** | 0.41**** | -28.21 (-4.70) | -30.43 (-5.07) |
| | Medial Orbitofrontal | 0.54**** | 0.48**** | -39.00 (-6.50) | -29.67 (-4.95) |
| | Paracentral | 0.28**** | 0.12*** | -38.81 (-6.47) | -21.92 (-3.65) |
| | Pars Opercularis | 0.48**** | 0.48**** | -34.67 (-5.78) | -38.25 (-6.38) |
| | Precentral | 0.39**** | 0.41**** | -35.08 (-5.85) | -38.81 (-6.47) |
| | Superior Frontal | 0.45**** | 0.41**** | -34.94 (-5.82) | -32.70 (-5.45) |
| Cingulate | Caudal Anterior Cing. | 0.54**** | 0.43**** | -47.43 (-7.91) | -37.15 (-6.19) |
| | Rostral Anterior Cing. | 0.57**** | 0.44**** | -42.29 (-7.05) | -41.26 (-6.88) |
| | Posterior Cing. | 0.44**** | 0.38**** | -31.92 (-5.32) | -28.46 (-4.74) |
| | Isthmus Cing. | 0.28**** | 0.18**** | -27.87 (-4.64) | -17.63 (-2.94) |
| Parietal | Postcentral | 0.40**** | 0.45**** | -44.41 (-7.40) | -52.00 (-8.67) |
| | Precuneus | 0.43**** | 0.39**** | -27.43 (-4.57) | -25.11 (-4.19) |
| | Superior Parietal | 0.29**** | 0.30**** | -31.58 (-5.26) | -32.19 (-5.36) |
| | Supramarginal | 0.39**** | 0.47**** | -31.37 (-5.23) | -37.93 (-6.32) |
| | Inferior Parietal | 0.21**** | 0.23**** | -20.22 (-3.37) | -22.35 (-3.72) |
| Occipital | Cuneus | 0.37**** | 0.32**** | -28.87 (-4.81) | -24.89 (-4.15) |
| | Lingual | 0.63**** | 0.40**** | -33.62 (-5.60) | -28.61 (-4.77) |
| | Lateral Occipital | 0.08ns | 0.15*** | -10.37 (-1.73) | -10.88 (-1.81) |
| | Pericalcarine | 0.13*** | 0.19**** | -15.45 (-2.57) | -19.65 (-3.28) |
| Temporal | Insula | 0.54**** | 0.63**** | -40.57 (-6.76) | -44.66 (-7.44) |
| | Transverse Temporal | 0.47**** | 0.41**** | -41.04 (-6.84) | -42.52 (-7.09) |
| | Parahippocampal | 0.42**** | 0.40**** | -45.01 (-7.50) | -35.31 (-5.89) |
| | Superior Temporal | 0.47**** | 0.49**** | -32.46 (-5.41) | -35.05 (-5.84) |
| | Entorhinal | 0.06ns | 0.03ns | -21.16 (-3.53) | -16.41 (-2.73) |
| | Banks of Sup. Temp. S. | 0.24**** | 0.20**** | -15.32 (-2.55) | -19.16 (-3.19) |
| | Fusiform | 0.50**** | 0.36**** | -24.14 (-4.02) | -21.10 (-3.52) |
| | Inferior Temporal | 0.30**** | 0.27**** | -18.45 (-3.08) | -18.82 (-3.14) |
| | Middle Temporal | 0.29**** | 0.39**** | -14.77 (-2.46) | -22.48 (-3.75) |
| | Termporal Pole | 0.33**** | 0.28**** | -27.16 (-4.53) | -27.74 (-4.62) |

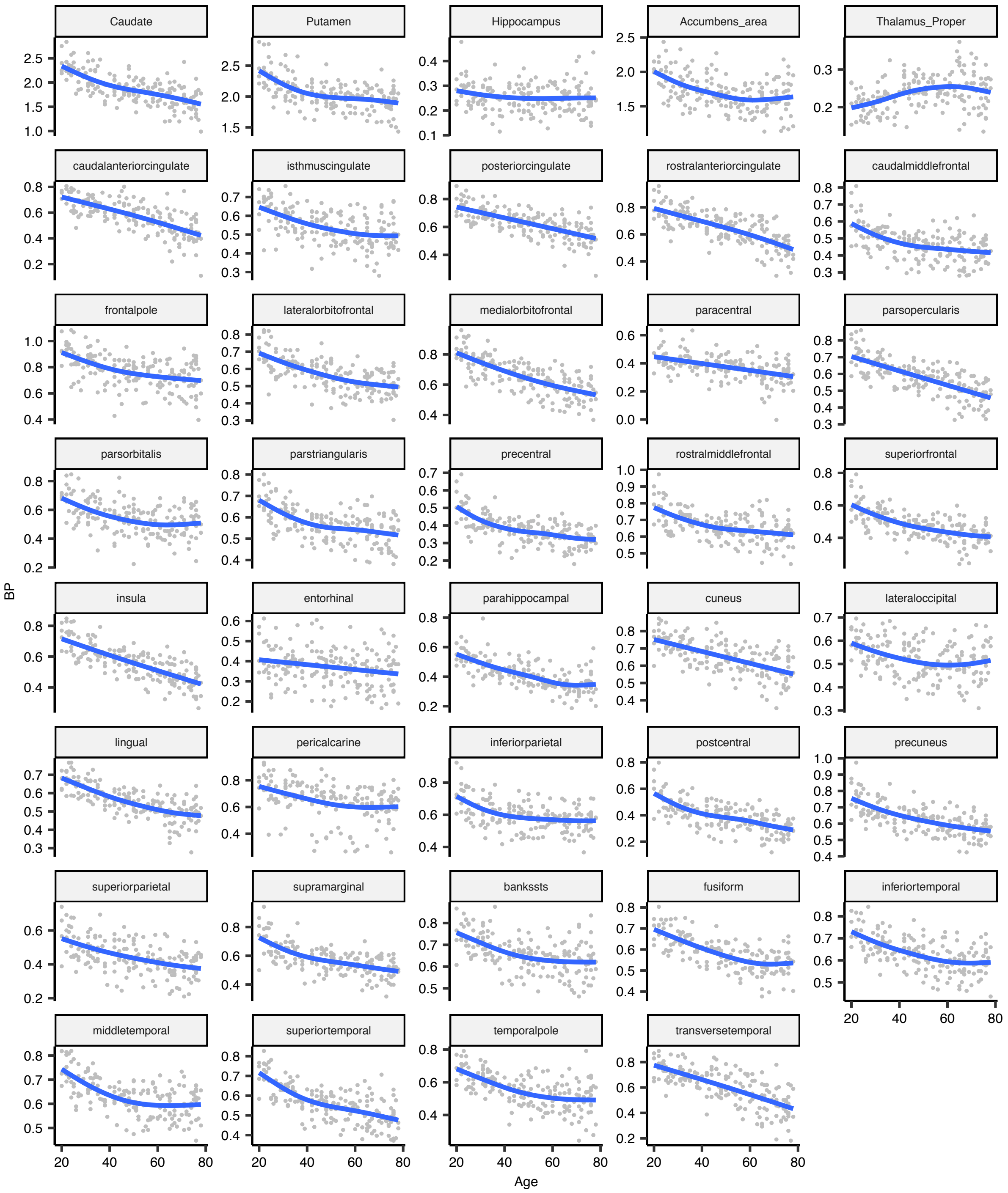


**Supplemental Figure 1:** Scatter plots of individual ROI-wise $BP_{ND}$ estimates in relation to age, and the corresponding GAM-predictions. Region-wise $BP_{ND}$ range (y-axis) was employed to emphasize interregional variations and similarities in age-related trajectories.

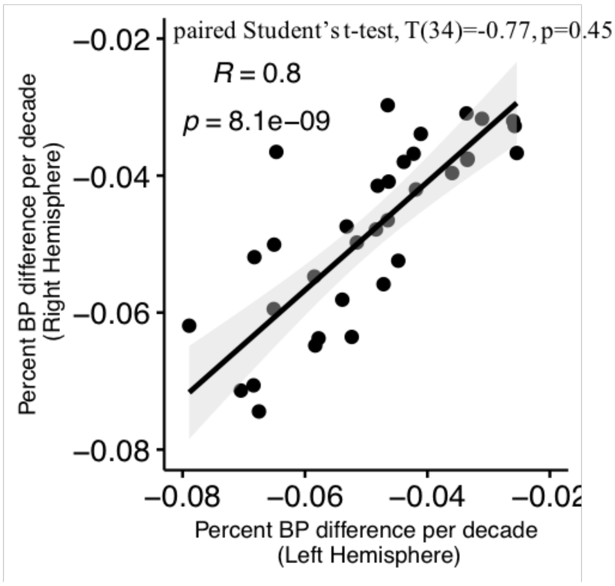


**Supplemental Figure 2:** ROI-based estimates of age-related SCH BP differences were not markedly different across the brain-hemispheres.

**Shape analysis of SCH BP age-trajectories.** ROI-based GAM-analysis suggested a high degree of commonalities across cortical age-trajectories (S.Fig 2). In order to reduce the number of cortical ROIs an automated shape-analysis to identify cortical regions in mutually similar relationship with age was conducted. To this end, hierarchical GAM-analysis including a common smooth factor of age (common age), regional smooth factors of age (regional age), regional intercepts, and subject-wise intercepts were first conducted incorporating all cortical ROIs in a multivariate fashion. This analysis confirmed the presence of a common age-factor, but evidence for significant regional differences across the cortical ROIs was also found (S.Table 2). Next, the common age-trend and regional intercepts were regressed out to orthogonalize regional age-patterns (see Main Fig 1A). These orthogonalized age-trajectories were next analysed using shape-respecting k-means clustering (REF) to find clusters of regions with mutually similar age-patterns. K-mean clustering identified three patterns of age-related differences: 1) the prefrontal cortex (PFC) together with posterior regions, 2) central regions, and 3) operculum together with the anterior cingulate cortex (ACC). The cluster incorporating DLPFC and posterior cortical regions was further subdivided into separate classes owing to their dopaminergic, topographic and functional disparities (Main Fig 1A). Assignment of Desikan-Killiany ROIs to these new ROIs are listed in Supplemental Table 3. An *Ad hoc* cluster-wise HGAM analysis confirmed internal coherence within each cortical cluster (Supplemental Table 4, cluster-age $p<10^{-16}$, region-age p = 0.01 – 1.0, Bonferroni-corrected p-values), indicating that interregional variations within clusters were non-significant, and therefore warranting the use of aggregate cortical regions-of-interest (n=4) in the subsequent analysis.

**Supplemental Table 2:** hierarchical GAM-statistics in cortical analysis (n=34 ROIs). Significant effects of smooth general age-trend and region-specific effects of age were observed.

| *term* | *edf* | *ref.df* | *F* | *p.value* |
|---|---|---|---|---|
| **Common Age** | 3 | 3 | 46.5 | **$<10^{-16}$** |
| **Regional Age** | 79 | 202 | 2921.4 | **<0.0001** |

**Supplemental Table 3**: Assignment of Desikan-Killiany ROIs (n=34) to the data-driven clusters (n=4) with similar age-based D1DR trajectories.

| | Region | FreeSurfer name |
|---|---|---|
| PFC | Caudal Middle Frontal | caudalmiddlefrontal |
| | Frontal Pole | frontalpole |
| | Pars Orbitalis | parsorbitalis |
| | Pars Triangularis | parstriangularis |
| | Rostral Middle Frontal | rostralmiddlefrontal |
| Operculum & ACC | Caudal Anterior Cing. | caudalanteriorcingulate |
| | Insula | insula |
| | Rostral Anterior Cing. | rostralanteriorcingulate |
| | Transverse Temporal | transversetemporal |
| Central | Cuneus | cuneus |
| | Lateral Orbitofrontal | lateralorbitofrontal |
| | Lingual | lingual |
| | Medial Orbitofrontal | medialorbitofrontal |
| | Paracentral | paracentral |
| | Parahippocampal | parahippocampal |
| | Pars Opercularis | parsopercularis |
| | Postcentral | postcentral |
| | Posterior Cing. | posteriorcingulate |
| | Precentral | precentral |
| | Precuneus | precuneus |
| | Superior Frontal | superiorfrontal |
| | Superior Parietal | superiorparietal |
| | Superior Temporal | superiortemporal |
| | Supramarginal | supramarginal |
| Posterior | Banks of Sup. Temp. S. | bankssts |
| | Entorhinal | entorhinal |
| | Fusiform | fusiform |
| | Inferior Parietal | inferiorparietal |
| | Inferior Temporal | inferiortemporal |
| | Isthmus Cing. | isthmuscingulate |
| | Lateral Occipital | lateraloccipital |
| | Middle Temporal | middletemporal |
| | Pericalcarine | pericalcarine |
| | Termporal Pole | temporalpole |

**Supplemental Table 4:** The data-driven classification was validated using cluster-wise HGAM to test for the common cluster and region-specific effects (Bonferroni-corrected p-values). All clusters showed a strong cluster age-effect and suppression of regional effects, corroborating cluster-wise similarity of age-effects.

| **Region** | ***p (Common Age)*** | ***p (Regional Age)*** |
|---|---|---|
| DLPFC | $<10^{-16}$ | 0.36 |
| Operculum & ACC | $<10^{-16}$ | 1.00 |
| Central | $<10^{-16}$ | 0.08 |
| Posterior | $<10^{-16}$ | **0.01** |

### Modulators of SCH BP differences.

**Supplemental Table 5:** Mixed linear effect model comparison statistics. D1DR availability served as the dependent variable and age was modeled as a fixed effect. Base model #1 included a common (fixed) effect of age and regional (random) effect of intercept. Base model #2 was an augmentation of model #1 with regional (random) effect of age (random slope). Inclusion of region-specific (random) effect of log-transformed white matter lesion volume abolished the effect of region-specific slope with age. Analysis was conducted on participants at age 40y and older (n = 116 / 109).

| Model | df | AIC | BIC | logLik | Test | L.Ratio | p-value |
|---|---|---|---|---|---|---|---|
| *Base model* | | | | | | | |
| #1 | 4 | -28.02 | -14.27 | 18.01 | | | |
| #2 | 6 | -31.02 | -10.39 | 21.51 | 1 vs 2 | 7.00 | 0.03 |
| *With region-specific effect of white matter lesion volume* | | | | | | | |
| #1 | 7 | -37.29 | -13.67 | 25.65 | | | |
| #2 | 9 | -33.37 | -2.99 | 25.68 | 1 vs 2 | 0.08 | 0.96 |

***Distinct associations between caudate D1DR and memory***

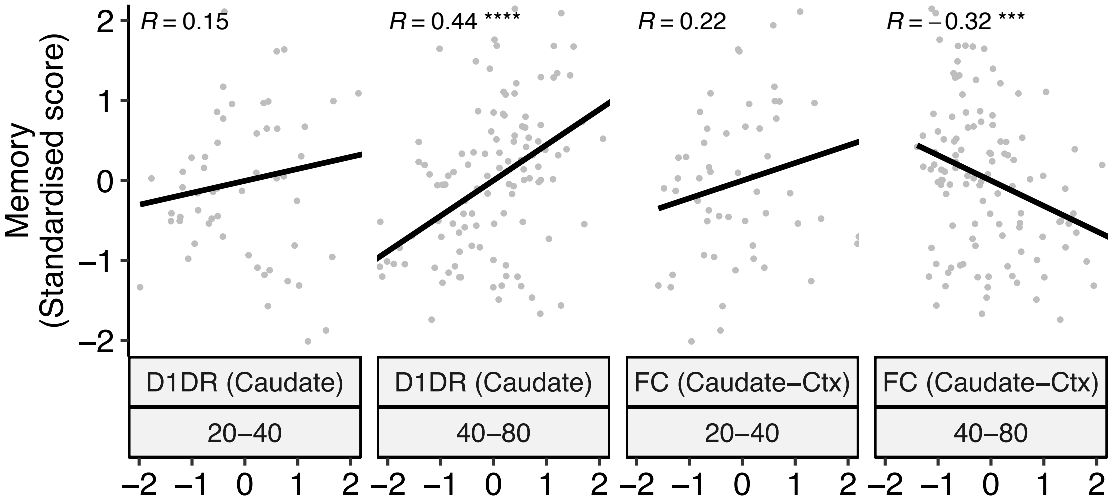


**Supplemental Figure 3:** FC-memory, but not D1DR-memory association was heavily modulated by age-segment.
